# The python-derived 16α-hydroxylated bile acid, pythocholic acid decreases food intake and increases jejunal fatty acid ethanolamides in mice

**DOI:** 10.1101/2022.12.01.518764

**Authors:** Sei Higuchi, Courtney Wood, Nicholas V. DiPatrizio, Akira Kawamura, Rebecca A. Haeusler

**Affiliations:** Department of Pharmaceutical Sciences, College of Pharmacy and Health Sciences, St. John’s University, Queens, NY 11439, USA; Naomi Berrie Diabetes Center and Department of Pathology and Cell Biology, Columbia University, New York, NY, USA; Division of Biomedical Sciences, School of Medicine, University of California, Riverside, CA, USA; Department of Chemistry, Hunter College of CUNY, New York, NY, USA

**Keywords:** Bile acids, pythocholic acid, OEA, NAPE-PLD, gastric emptying

## Abstract

**Objective:** Modulation of bile acid (BA) structure is a potential strategy for obesity and metabolic disease treatment. BAs act not only as signaling molecules involved in energy expenditure and glucose homeostasis, but also as regulators of food intake. The structure of BAs, particularly the position of the hydroxyl groups of BAs impacts food intake partly by intestinal effects: (1) modulating the activity of N-acyl phosphatidylethanolamine phospholipase D (NAPE-PLD), which produces the anorexigenic bioactive lipid oleoylethanolamide (OEA), or (2) regulating lipid absorption and the gastric emptying-satiation pathway. We hypothesized that 16α-hydroxylated BAs uniquely regulate food intake, because of the long intermeal intervals in snake species in which these BAs are abundant. However, the effects of 16α-hydroxylated BAs in mammals are completely unknown, because 16α-hydroxylated BAs are not naturally found in mammals. To test the effect of 16α-hydroxylated BAs on food intake, we isolated the 16α-hydroxylated BA pythocholic acid from ball pythons (*Python regius*).

**Methods:** Pythocholic acid or deoxycholic acid (DCA) were given by oral gavage in mice. DCA is known to increase NAPE-PLD activity better than other mammalian BAs. We evaluated food intake, OEA levels and gastric emptying in mice.

**Results:** We successfully isolated pythocholic acid from ball pythons for experimental use. Pythocholic treatment significantly decreased food intake compared with DCA treatment, and this was associated with increased jejunal OEA, but no change in gastric emptying or lipid absorption.

**Conclusion:** The exogenous bile acid pythocholic acid is a novel regulator of food intake and the satiety signal OEA in the mouse intestine.

**Highlights:**

- Pythocholic acid decreases food intake.
- Pythocholic acid increases intestinal OEA and other fatty acid ethanolamides.
- The effects of pythocholic acid on OEA and hypophagia are greater than the effects of DCA.
- Pythocholic acid does not affect lipid absorption or gastric emptying.

## 1. INTRODUCTION

Bile acids (BAs) are synthesized from cholesterol in the liver and promote lipid absorption in the intestine. BAs act as signaling molecules by activating BA receptors, Takeda G protein-coupled receptor 5 (TGR5) and transcription factor farnesoid X receptor (FXR) [1; 2]. Activation of TGR5 increases energy expenditure and prevents diet-induced obesity [3; 4]. TGR5 signaling is also involved in insulin secretion [5] and GLP-1 secretion [6]. FXR activation mitigates insulin resistance in diabetic rodents by improving insulin signaling [7], GLP-1 secretion [8] and inhibiting gluconeogenic gene expression [9]. Recently, BAs were shown to act in the hypothalamus to stimulate satiety [4; 10] Thus, BA receptors may be a potential therapeutic target for obesity and metabolic diseases, because of their traits as signaling molecules involved in energy balance and glucose homeostasis.

In addition to the involvement of BA receptors in energy balance and glucose homeostasis, BAs can also regulate food intake via BA receptor-independent mechanisms. One is an intestinal lipid sensing mechanism. Our previous study indicated that BAs regulate gastric emptying and satiation by determining lipid access into the distal intestine and the distally-abundant fat sensing receptor, GPR119 [11]. A second mechanism is via BA-induced allosteric modulation of bioactive lipid synthesis. Magotti et al showed that BAs hydroxylated at carbon 3α and 12α bind and activate the enzyme N-acyl phosphatidylethanolamine phospholipase D (NAPE-PLD), an enzyme that produces fatty acid ethanolamides (FAEs) [12]. In line with this, we showed that oleoyl ethanolamide (OEA) concentrations were reduced in the jejunal epithelium of mice lacking the 12α-hydroxylated BAs, cholic acid (CA) and deoxycholic acid (DCA) [11]. Jejunal OEA is a bioactive signaling lipid that induces satiety–defined as an increase in the time interval between meals–via PPARα activation and the vagally-mediated gut-brain axis [13–16]. Thus the structure of BAs, particularly the position of the hydroxyl groups, determines the production of the satiety molecule OEA.

Clinical and animal studies have indicated that OEA is a prospective therapeutic target for obesity and metabolic diseases. A human study shows that a single nucleotide polymorphism of NAPE-PLD is associated with severe obesity [17]. OEA treatment decreases body weight and fat mass by reducing the desire to eat and appetite for sweet foods in obese subjects [18; 19]. Also, preclinical studies have established the physiological role of OEA. Intestinal OEA induces dopamine release in the brain to reduce fat appetite in mice [20]. OEA treatment reduces total food intake and meal size in diet-induced obese animal models as well as a mouse model of Prader-Willi syndrome, which is a genetic disorder that causes childhood-onset hyperphagia and obesity [21; 22]. The mechanism of OEA-induced satiety and hypophagia is suggested to be mediated by PPARα activation in the intestine [14; 15]. However, some adverse effects of systemic PPARα activation have been reported such as hepatocellular carcinoma or chemotaxis [23–25]. Overexpression of PPARα induces lipid accumulation in cardiomyocytes [26; 27] and glucose intolerance in skeletal muscle [28]. Thus, locally increasing intestinal OEA presents an attractive alternative against obesity, because it may mitigate the side effects of systemic PPARα activation.

We focused on the atypical research animal, pythons, which have a unique system to promote and sustain high levels of OEA in the intestine. In pythons, jejunal OEA is increased 300-fold and sustained for at least 2 days after a meal [29]. For comparison, jejunal OEA levels in rodents are increased 2-10 fold after a meal, and are reduced to pre-meal levels with in a few hours in rodents [11; 15; 30; 31]. Pythons consume a large meal at infrequent intervals (a couple of months or a year). The dramatic increase in postprandial OEA in pythons is considered to be a potential mechanism contributing to pythons’ long intermeal interval [32]. However, the regulation of intestinal OEA production in pythons is incompletely defined.

In this work, we examined the role of the unique BA in pythons, pythocholic acid, in the production of OEA. Pythocholic acid was discovered in 1950 by Haslewood and Wootton as a major BA in pythons [33; 34]. Pythocholic acid is a unique BA because of its structure, which has hydroxyl groups at positions 3α-, 12α- and 16α-[34]. Hydroxylation at 16α-is common in specific snakes (Cylindrophiidae, Uropeltidae, Boidae and Pythonidae) and birds (Shoebill), but is not seen in mammals[35]. Pythocholic acid has been found in specific snakes such as boas and pythons that possess long meal intervals [35]. We hypothesized that the unique structure of pythocholic acid promotes NAPE-PLD activity and increases intestinal OEA production [11; 12; 36; 37]. Here, we tested the effects of pythocholic acid on OEA production and satiety in mice.

## 2. MATERIALS AND METHODS

### 2.1.1. Animals

Wild-type mice are C57BL/6J (The Jackson Laboratory #000664). Mice were fed a normal chow diet (3.4 kcal/g, Purina 5053, 24.7% kcal from protein, 62.1% carbohydrate and 13.2% fat). Mice were provided with the diet and water ad libitum and maintained on a 12-hour light/dark cycle, set with lights on at 7 AM. All experiments were approved and conducted according to the guidance of the Columbia University Institutional Animal Care and Use Committee.

### 2.1.2. Food intake measurements

For food intake measurements, mice were individually housed, and the food dispenser was located inside the cage.

### 2.1.3. Gastric emptying

Solid gastric emptying was measured following ingestion of a normal chow diet as we previously reported [11]. Mice were fasted for 16-h with free access to water, then allowed access to chow diet for 1-h. Mice were food-deprived again for 2-h before euthanasia. Food intake during the 1-h feeding period was measured using a food dispenser. Food content in the stomach was measured at euthanasia. Solid gastric emptying was calculated by the following formula: solid gastric emptying (%) = {1-(food content in stomach/food intake)}x100. For python bile or BA treatment, mice were orally gavaged with 20 mg/kg of these BAs in 1.5% NaHCO_3_ or equivalent dose of pythocholic acid-containing python bile for 2 or 4 consecutive days at 6 PM.

### 2.2. Extraction of pythocholic acid from python bile

Euthanized ball pythons (Python regius) were donated by Mr. Dustin Leahy (Piedthonidae Exotics). Bile was collected from the gall bladder. Pythocholic acid and Tauro-conjugated pythocholic acid were purified using HyperSep C8 solid phase extraction (SPE) cartridge (Thermo Scientific, Waltham, MA, Cat # 60108-309) and silica gel chromatography (Figure 4 A). Python bile (0.5 mL) was suspended in 2 mL of aqueous HCl (0.01 M). The acidified suspension was extracted with ethyl acetate (2 mL × 5). The ethyl acetate extract was subjected to liquid chromatography/mass spectrometry (LC/MS) on Agilent iFunnel 6550 Q-ToF LC/MS System. The total ion chromatogram (TIC) in the positive ion mode showed a major peak with *m/z* 431.2769 (Figure 4 B and C), which was consistent with the sodium adduct of pythocholic acid [M+Na]^+^ (calculated *m/z* 431.2765). The ethyl acetate extract was then subjected to further purification with SPE cartridge, from which the major constituent was eluted with 20% methanol in water. NMR of the purified material (4 mg, clear solid) confirmed its identity as pythocholic acid [38]. The aqueous layer from the ethyl acetate extraction was also subjected to LC/MS to give a predominant peak with *m/z* 516.2999 (Figure 4 D and E), which corresponded to the protonated form of Tauro-conjugated pythocholic acid [M+H]^+^ (calculated *m/z* 516.2989). The major compound was purified with silica gel chromatography to give clear solid (45.5 mg, R_*f*_ 0.2, 30% methanol in dichloromethane). The steroidal moiety of the purified material was confirmed to be pythocholic acid after deconjugation in base [39].

**Figure 1.**
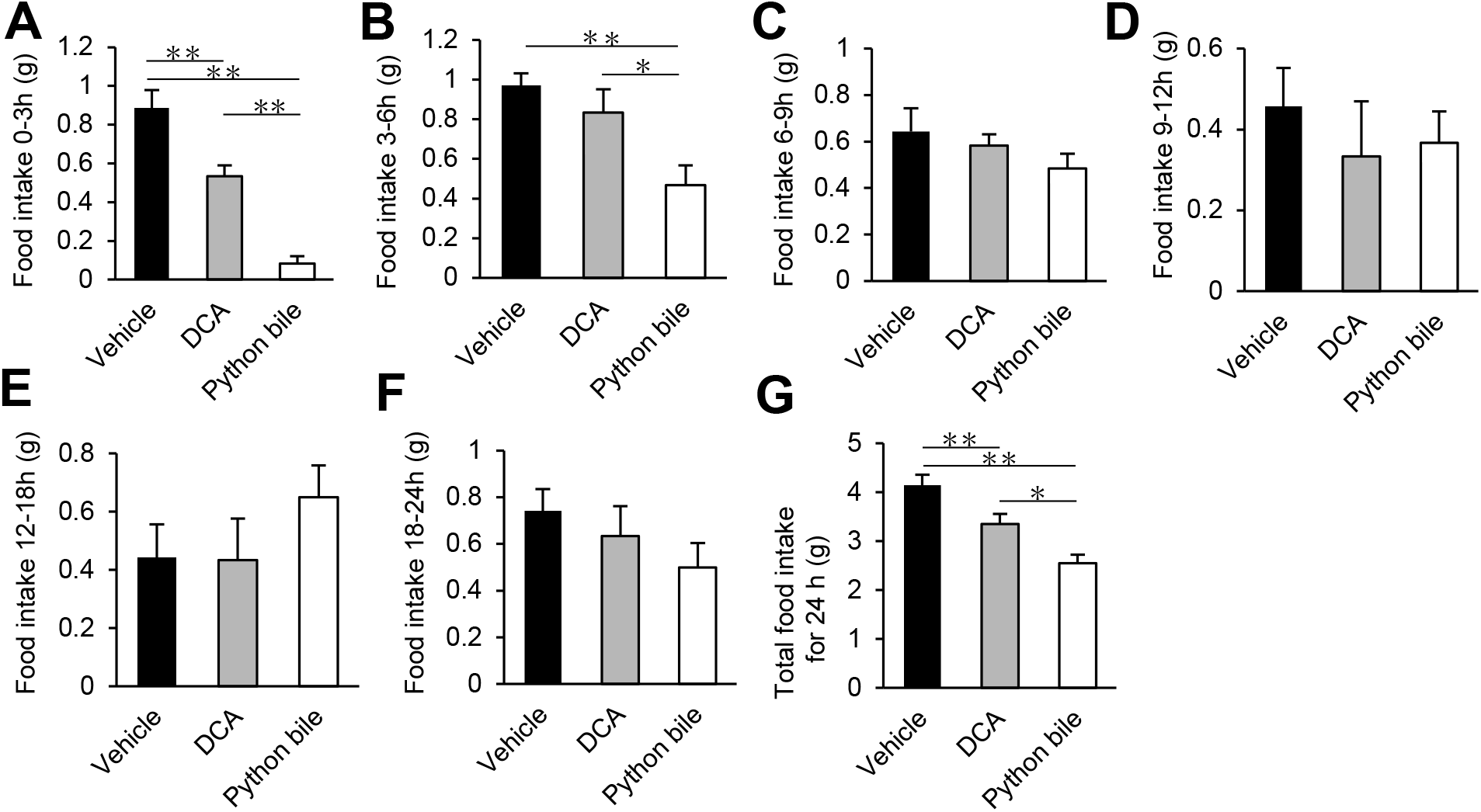
Ad libitum food intake after single dose administration of deoxycholic acid (DCA) or python bile at 6 pm. Food intake after treatment 0-3 hours (A), 3-6 hours (B), 6-9 hours (C), 9-12 hours(D), 12-18 hours (E), 18-24 hours (F), and total food intake for 24 hours (G) (n= 6-7 for each group). \**p*<0.05, \*\**p*<0.01 (one-way repeated ANOVA).

**Figure 2.**
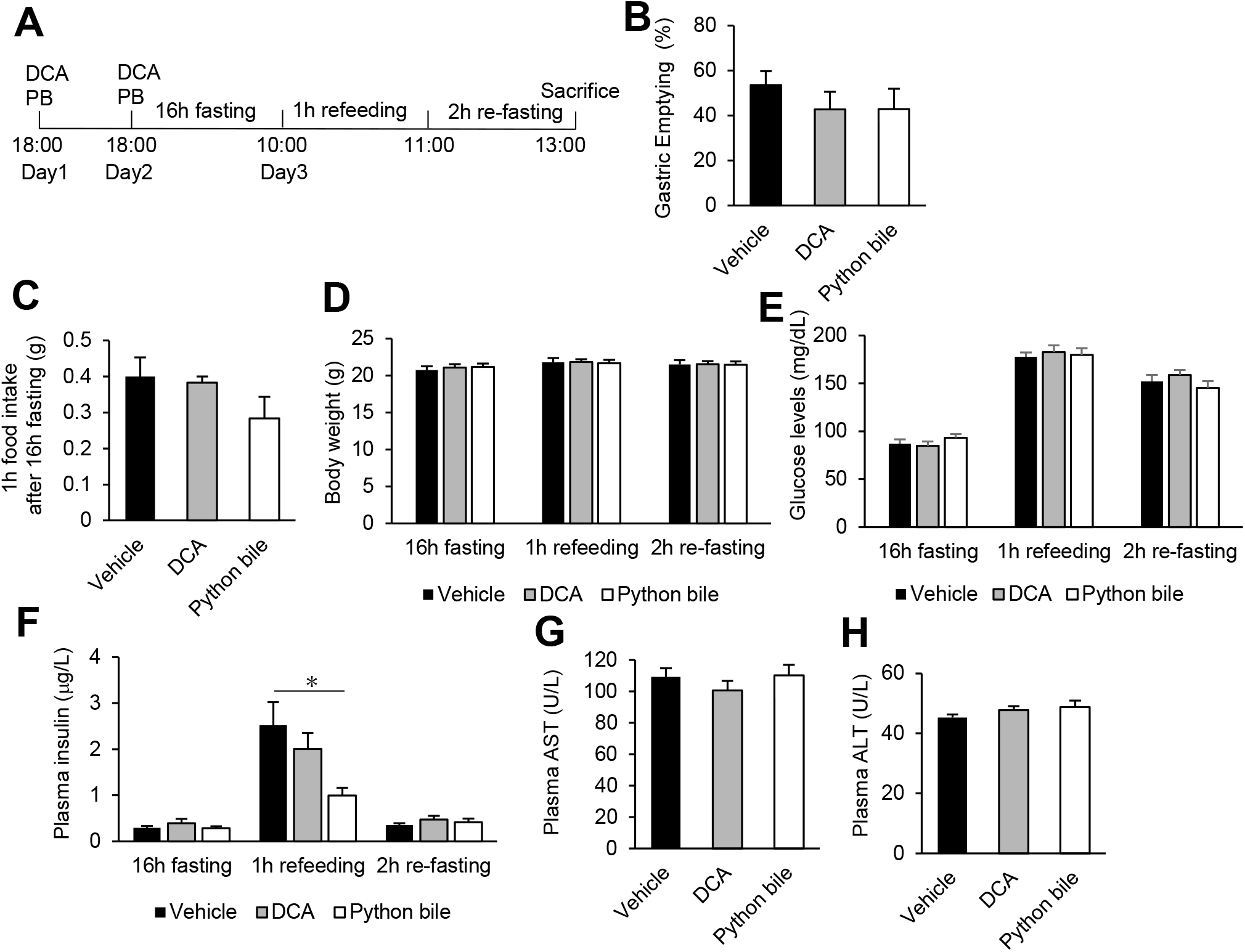
Effects of DCA and python bile on gastric emptying, blood glucose and insulin levels, after two days of once-daily treatment. (A) experimental schedule, (B) gastric emptying, (C) 1 h refed food intake after fasting, (D) body weight, (E) blood glucose levels, (F) plasma insulin levels, (G) plasma AST levels and (H) plasma ALT levels. (n= 6-7 for each group). \**p*<0.05 (one-way repeated ANOVA).

**Figure 3.**
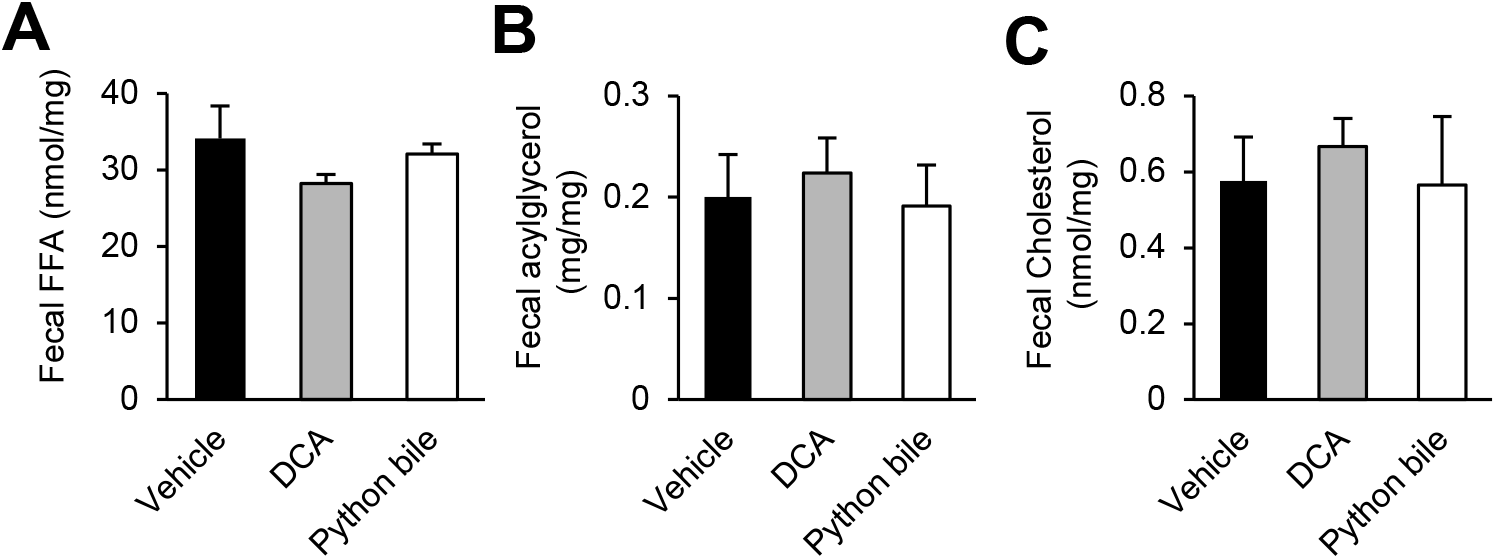
Influence of DCA and python bile on lipid absorption. (A) fecal free fatty acid, (B) fecal acylglycerol and (C) fecal cholesterol levels. Feces were collected for 24 before the gastric emptying experiment (n= 6-7 for each group).

**Figure 4.**
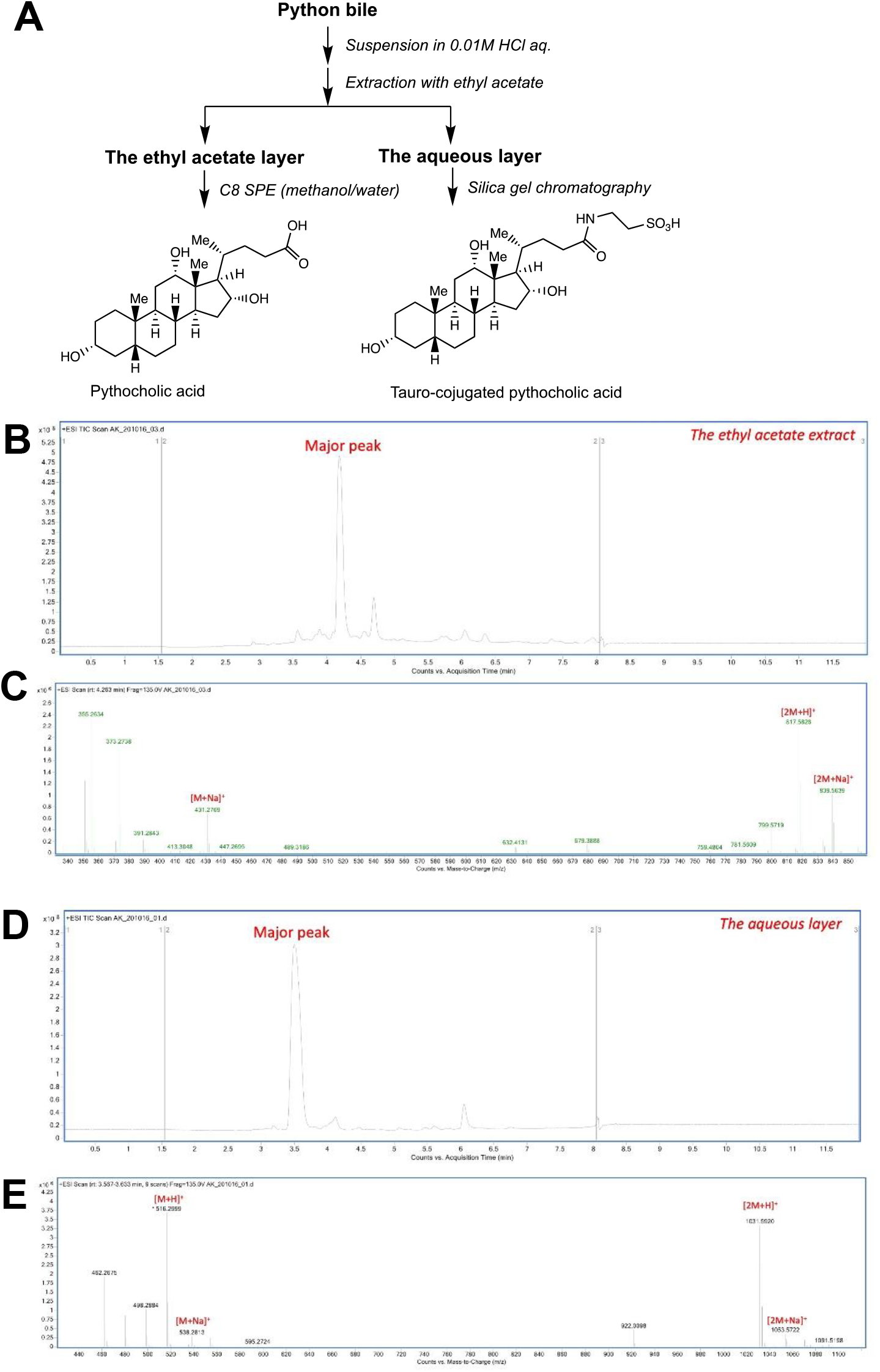
Purification of pythocholic acid and its taurine conjugate from python bile. (A) Python bile was first fractionated by partition between organic (ethyl acetate) and aqueous phases. The ethyl acetate phase was further purified by C8 solid phase extraction to give pythocholic acid, whereas the aqueous phase was subjected to silica gel chromatography to give Tauro-conjugate. (B-C) Liquid Chromatography/Mass Spectrometry (LC/MS) analysis of the ethyl acetate extract from python bile. (B) Total ion chromatogram (TIC) in the positive mode. The major peak was observed around 4.2-4.3 min. (C); High resolution electrospray mass spectrometry (HR ESI MS) of the major peak (4.2-4.3 min). The MS signal at *m/z* 431.2769 was consistent with the sodium adduct of pythocholic acid [M+Na]^+^ (calculated *m/z* 431.2765). (D-E) Liquid Chromatography/Mass Spectrometry (LC/MS) analysis of the aqueous phase from python bile (after ethyl acetate extraction). (D) Total ion chromatogram (TIC) in the positive mode. The major peak was observed around 3.5 min. (E) High resolution electrospray mass spectrometry (HR ESI MS) of the major peak (~3.5 min). The MS signal at *m/z* 516.2999 was consistent with the protonated form of Tauro-conjugated pythocholic acid [M+H]^+^ (calculated *m/z* 516.2989).

### 2.3. Blood and plasma analysis

Blood glucose was measured in mice by tail vein bleeding using a OneTouch glucose monitor and strips (LifeScan). Plasma was obtained after centrifuging blood collected in EDTA-coated tubes for 15 mins at 2,500 g. Plasma insulin was measured using an ELISA kit (# 10-1247-01, Mercodia) according to the manufacturer’s protocol. Plasma alanine aminotransferase (ALT) (MAK052, Sigma) and aspartate aminotransferase (AST) (MAK055, Sigma) were measured using dedicated kits according to the manufacturer’s protocol

### 2.4. Fecal lipid extraction and measurement

Feces for 24 hours was collected from each cage after 2 or 4 consecutive days of treatment of BA or python bile. Collected feces were dried in an incubator for 16h at 42 °C and 100 mg of feces were homogenized with 1 ml of 1M NaCl. One ml of homogenized solution was added into 6 ml of chloroform:methanol (2:1). Chloroform layers were collected after centrifugation, then evaporated under N2 flow until dry. One ml of 2% Triton X-100 in chloroform was added and evaporated again under N2 flow. One ml of ddH2O was added and vortexed until sample dissolved. Fecal free fatty acid (FFA, Wako; HR Series NEFA-HA(2),), triglycerides (Infinity; Thermo Scientific) and cholesterol (Cholesterol E, Wako Diagnostics) content were assessed using a colorimetric assay.

### 2.5. Intestinal lipid extraction and analysis of FAEs and MAGs

Lipid extraction and analysis were performed as previously described [40–42] Frozen tissue of mucosal intestine and mediobasal hypothalamus was blade-homogenized in 1.0 mL of methanol solution containing the internal standards, [^2^H_5_] 2-arachidonylglycerol (2-AG), [^2^H_4_]-anandamide (AEA), [^2^H_4_]-OEA (Cayman Chemical, Ann Arbor, MI, USA). Lipids were extracted with chloroform (2 mL) and washed with water (1 mL). Organic (lower) phases were collected and separated by open-bed silica gel column chromatography as previously described [43]. Eluate was gently dried under N2 stream (99.998% pure) and resuspended in 0.2 mL of methanol:chloroform (9:1), with 1 μL injection for ultra-performance liquid chromatography/tandem mass spectrometry (UPLC/MS/MS) analysis.

Data acquisition was performed using an Acquity I Class UPLC with in-line connection to a Xevo TQ-S Micro Triple Quadrapole Mass Spectrometer (Waters Corporation, Milford, MA, USA) and accompanying electrospray ionization (ESI) sample delivery. Lipids were separated using an Acquity UPLC BEH C18 column (2.1 × 50 mm i.d., 1.7 μm, Waters) and inline guard column (UPLC BEH C_18_ VanGuard PreColumn; 2.1 × 5 mm i.d.; 1.7 μm, Waters). Lipids were eluted by a gradient of water and methanol (containing 0.25% acetic acid, 5 mM ammonium acetate) at a flow rate of 0.4 mL/min and gradient: 80% methanol 0.5 min, 80% to 100% methanol 0.5 to 2.5 mins, 100% methanol 2.5 to 3.0 mins, 100% to 80% methanol 3.0 to 3.1 mins, and 80% methanol 3.1 to 4.5 mins. Column was maintained at 40°C and samples were kept at 10°C in sample manager. MS detection was in positive ion mode with capillary voltage maintained at 1.10 kV and Argon (99.998%) was used as collision gas. Cone voltages and collision energies for respective analytes: AEA = 30v, 14v; OEA = 28v, 16v; DHEA = 30v, 16v; 2-AG (20:4) = 30v, 12v 2-OG (18:1) = 42v, 10v; 2-DG (22:6)= 34v, 14v; 2-LG (18:2) = 30v, 10v; [^2^H_4_]-AEA = 26v, 16v; [^2^H_4_]-OEA = 48v, 14v; [^2^H_5_]-2-AG = 25v, 44v. Lipids were quantified using a stable isotope dilution method detecting proton or sodium adducts of the molecular ions [M + H/Na]^+^ in multiple reaction monitoring (MRM) mode. For many MAGs, acyl migration from sn-2 to sn-1 glycerol positions is known to occur; for these analytes, the sum of these isoforms is presented. Tissue processing and LCMS analyses for experiments occurred independently of other experiments. Extracted ion chromatograms for MRM transitions were used to quantitate analytes: AEA (*m/z* = 348.3 > 62.0), OEA (*m/z* = 326.4 > 62.1), DHEA, (*m/z* = 372.3 > 62.0), 2-AG (*m/z* = 379.3 > 287.3), 2-OG (m/z = 357.4>265.2), 2-DG (*m/z* = 403.3 > 311.2), and 2-LG (m/z = 355.3>263.3), with [^2^H_4_]-AEA (*m/z* = 352.4 > 66.1) as internal standard for AEA and DHEA, [^2^H_4_]-OEA (*m/z* = 330.4 > 66.0) as internal standard for OEA, and [^2^H_5_]-2-AG (*m/z* = 384.3 > 93.4) as internal standard for 2-AG, 2-OG, 2-DG, and 2-LG. One “blank” sample was processed and analyzed in the same manner as all samples, except no tissue was included. This control revealed no detectable endocannabinoids and related lipids included in our analysis.

### 2.6. Statistics

Results are presented as mean ± SEM. Data were analyzed by one- and two-way ANOVA with Tukey’s multiple comparisons test or Student’s t-test.

## 3. RESULTS

### 3.1. Python bile treatment reduced food intake independent of slow gastric emptying and lipid absorption

Pythocholic acid is not commercially available. Thus, we first used python bile to test the effect of pythocholic acid on food intake because the major bile acid (over 70%) in pythons is reported to be pythocholic acid [34]. We measured pythocholic acid in the bile of ball pythons by LC-MS/MS and found 6 mg/mL of pythocholic acid in the bile of ball pythons. We treated wild-type mice with a single dose of vehicle or python bile (100 μL, which is estimated to yield a dose of approximately 20 mg/kg of pythocholic acid) by oral gavage for food intake measurement. As a positive control, we treated a parallel set of mice with DCA, because kinetic experiments demonstrate that DCA increases NAPE-PLD activity better than other BAs [37]. Mice were fed ad libitum prior to the gavage, which was delivered at 6 pm. We measured food intake every 3 or 6 hours. DCA decreased food intake in the first 3 hours after treatment (Figure 1A), but subsequent food intake returned to normal (Figure 1B-F). On the other hand, python bile decreased food intake more profoundly (Figure 1A-B), and this was sustained over six hours before returning to normal (Figure 1C-F). Python bile strongly decreased cumulative food intake for 24h relative to DCA treatment (Figure 1G). We then examined putative mechanisms underlying the hypophagic effect of python bile in mice.

Lipids in the distal small intestine suppress food intake by slowing gastric emptying and inducing satiation [44–47]. Previously, we reported that modulation of bile acid composition by Cyp8b1 deficiency caused slow gastric emptying because of impaired lipid absorption [11]. To evaluate the effect of python bile on gastric emptying and lipid absorption, we gave DCA or python bile once a day for two days before the gastric emptying experiment. After 2 days of treatment of DCA or python bile, we fasted mice for 16-h, then allowed them to access chow for 1-h, then removed food for 2-h (Figure 2A). We measured individual food intake and stomach contents to determine the percentage of food eaten that had emptied from the stomach. There was no significant difference in gastric emptying after DCA or python bile treatment (Figure 2B). We determined that “refed” food intake was not different during the 1h feeding period after 16h fasting (Figure 2C). Also, DCA and python bile did not affect body weight or glucose levels during the experiment (Figure 2D-E). Python bile decreased plasma insulin levels after 1h refeeding compared with the vehicle treatment group (Figure 2F). But there were no differences in plasma insulin levels after 16h fasting and 2h refasting between vehicle, DCA and python bile treated groups (Figure 2F). At the end of the experiment, we observed no differences in plasma AST and ALT levels due to DCA or python bile treatment, suggesting that oral gavage of python bile at hypophagic doses is not toxic for mice (Figure 2G-H).

It is well-known that bile acids promote lipid absorption, and the effects of BA composition on lipid absorption can contribute to systemic energy homeostasis [11; 48–51]. To evaluate the lipid absorption after 1 days of DCA and python bile treatment, we measured fecal excretion of free fatty acids (FFA), acylglycerols and cholesterol. We did not find a significant difference in fecal FFA, acylglycerol or cholesterol after DCA and python bile treatment (Figure. 3 A-C). These data implied that python bile induced a hypophagic effect that was independent of gastric emptying and lipid absorption.

### 3.2. Extraction and purification of pythocholic acid from python bile

Bile in python contains not only bile acids but also other lipids and biliary contents. To specifically test the effects of pythocholic acid on food intake, we extracted and purified pythocholic acid from ball python bile. Python bile was first acidified to fully protonate pythocholic acid, which made it easier to extract with ethyl acetate (Figure 4A). The tauro-conjugate remained in the aqueous phase during the extraction due to its polar taurine moiety. LC/MS analyses revealed molecular ions that are consistent with pythocholic acid and tauro-conjugate in the ethyl acetate extract and the aqueous phase, respectively (Figure 4B-E). The ethyl acetate extract was further purified by reversed phase C8 SPE to give pythocholic acid (4 mg), whereas the aqueous phase was subjected to silica gel chromatography to obtain tauro-conjugated pythocholic acid (45.5 mg).

### 3.3. Pythocholic acid treatment is sufficient to reduce food intake and does not influence gastric emptying or lipid absorption

To test the effect of pythocholic acid on food intake, we gave a single dose of pythocholic acid (20 mg/kg) by oral gavage to ad libitum fed mice. DCA (20 mg/kg) was given to mice as a positive control. Pythocholic acid and DCA were delivered at 6 pm of the test day, and food intake was measured every 3 or 6 hours. Both DCA and pythocholic acid decreased food intake in the first three hours after treatment (Figure 5A). Food intake returned to normal in all groups between 3 and 18 hours (Figure 5B-E). Unexpectedly, DCA and pythocholic treatment decreased food intake again between 18 and 24 hours, and the hypophagic effect of pythocholic acid was stronger than that of DCA, though this was a period of low food intake in all groups (Figure 5F). Cumulatively, pythocholic acid significantly decreased food intake over 24 hours compared with vehicle or DCA (Figure 5G).

**Figure 5.**
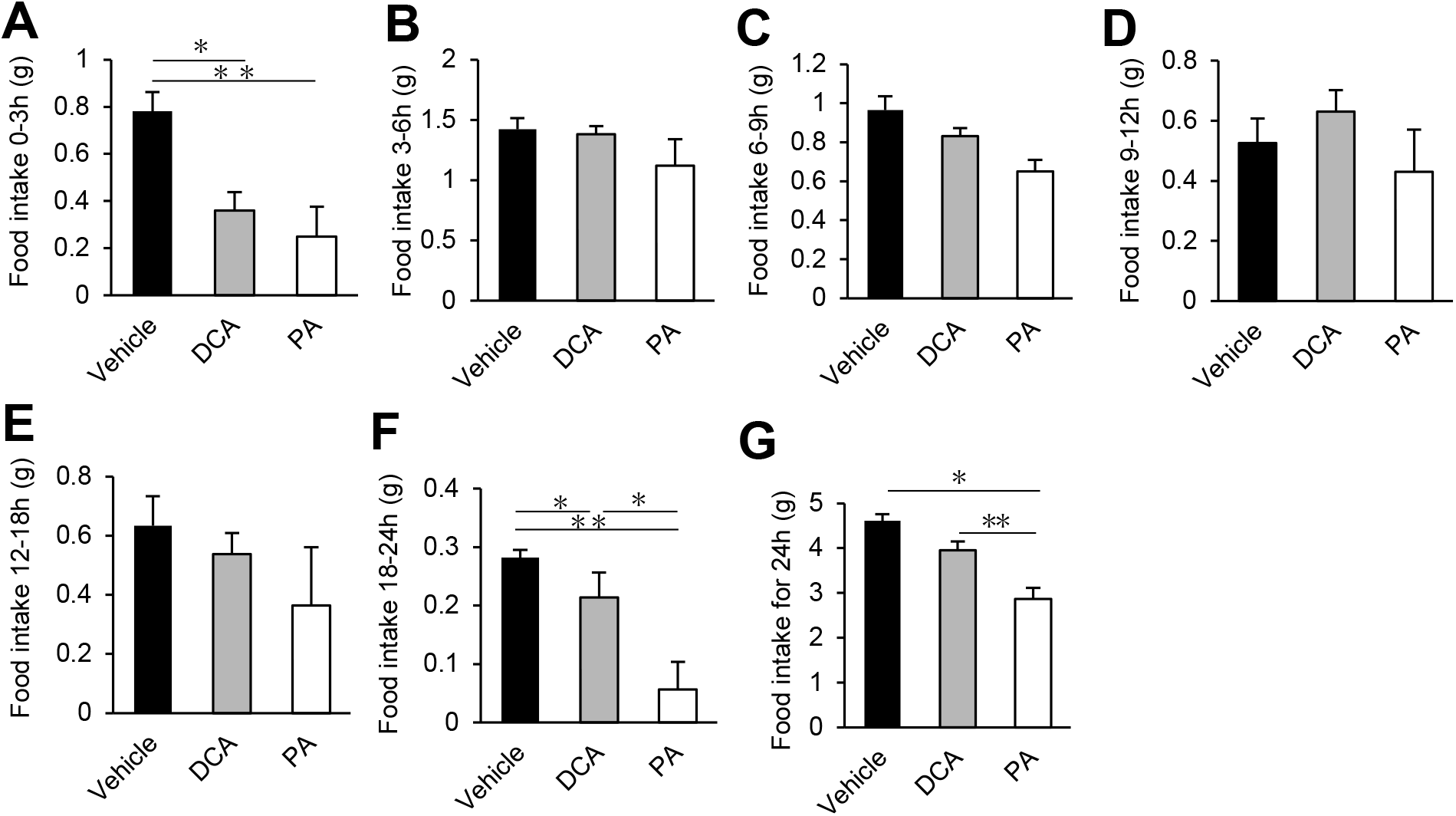
Ad libitum food intake after single dose administration of DCA or pythocholic acid (PA) at 6 pm. Food intake after treatment 0-3 hours (A), 3-6 hours (B), 6-9 hours (C), 9-12 hours(D), 12-18 hours (E), 18-24 hours (F), and total food intake for 24 hours (G) (Vehicle; n=5, DCA; n=5, PA; n=3). \**p*<0.05, \*\**p*<0.01 (one-way repeated ANOVA).

To evaluate the effect of pythocholic acid on gastric emptying, DCA or pythocholic acid were given by oral gavage for 2 days (Figure 6A). There were no significant differences in gastric emptying after DCA or pythocholic acid treatment in mice (Figure 6B). Food intake after overnight fasting was not different during the 1h refeeding period between any groups (Figure 6C), and there were no differences in body weight or insulin levels during the experiment (Figure 6D&F). Pythocholic acid increased the blood glucose levels after 1h refeeding compared with vehicle treatment group (Figure 6E), but there were no differences in blood glucose between groups at other time points. At the end of the experiment, there were no differences in plasma AST and ALT levels between groups (Figure 6H). We also did not find any differences in fecal FFA, acylglycerol, or cholesterol levels between DCA and pythocholic acid-treated groups (Figure 7A-C). Thus, we concluded that pythocholic acid is sufficient to induce hypophagia independent of gastric emptying and lipid absorption.

**Figure 6.**
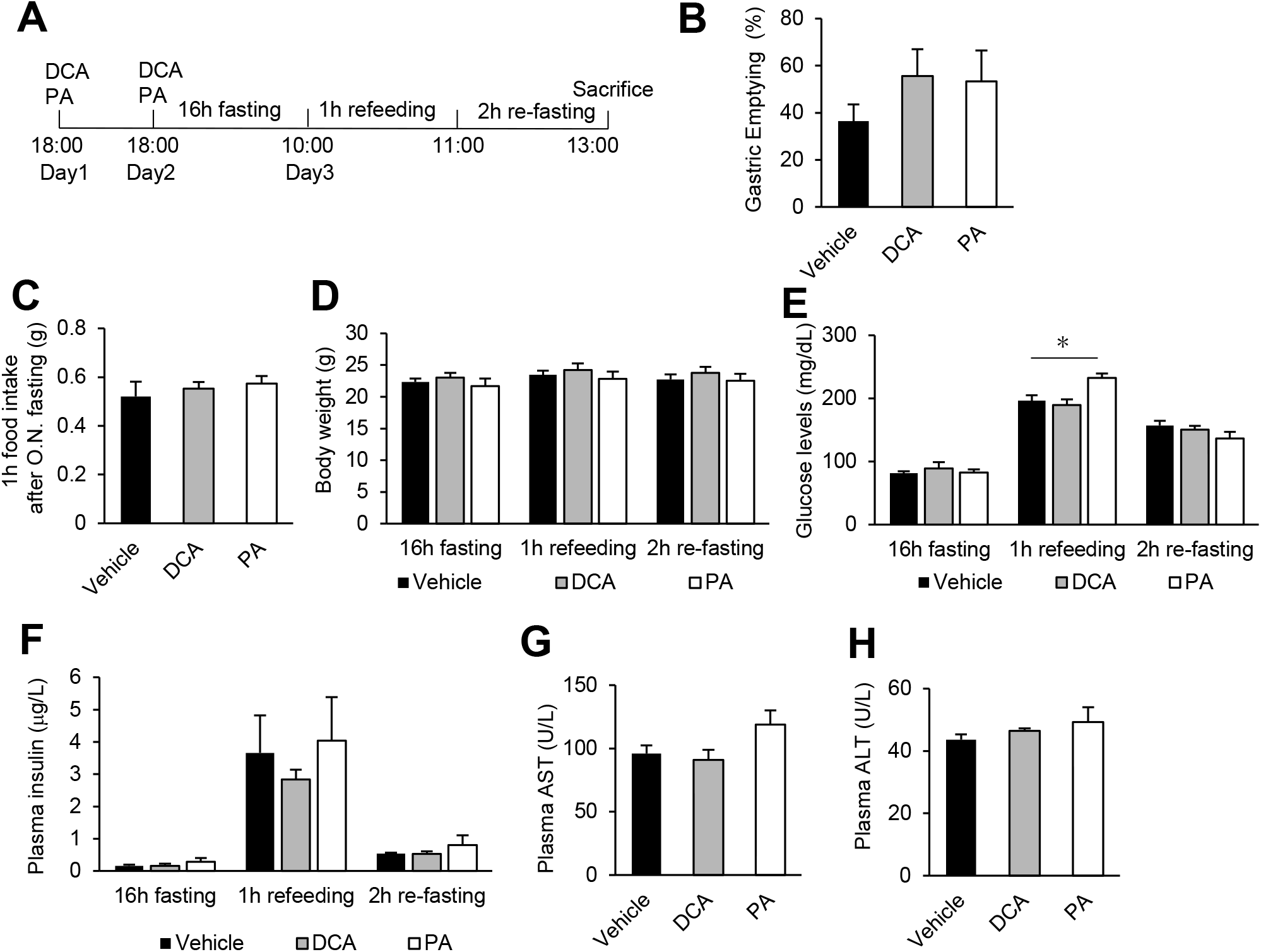
Effects of DCA and PA on gastric emptying, blood glucose and insulin levels, after two days of once-daily treatment. (A) experimental schedule, (B) gastric emptying, (C) 1 h refed food intake after fasting, (D) body weight, (E) blood glucose levels, (F) plasma insulin levels, (G) plasma AST levels and (H) plasma ALT levels. (Vehicle; n=5, DCA; n=5, PA; n=3). \**p*<0.05 (one-way repeated ANOVA).

**Figure 7.**
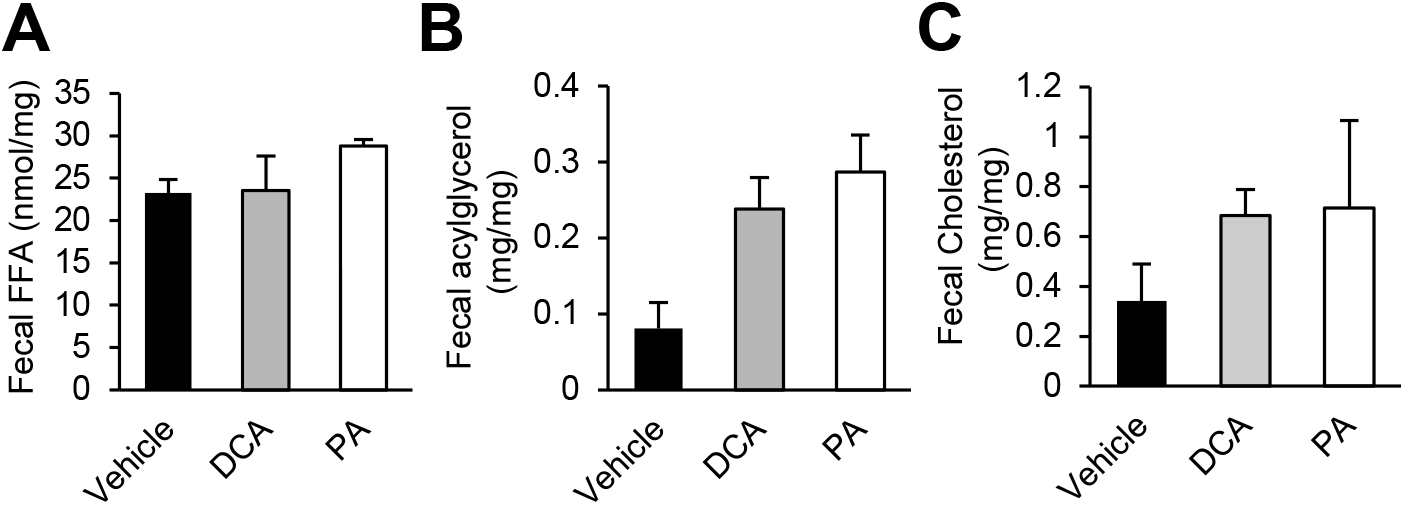
Influence of DCA and PA on lipid absorption. (A) fecal free fatty acid, (B) fecal acylglycerol and (C) fecal cholesterol levels. Feces were collected for 24 before the gastric emptying experiment. (Vehicle; n=5, DCA; n=5, PA; n=3).

### 3.4. Pythocholic acid promoted jejunal NAPE-PLD dependent fatty acid ethanolamide synthesis

Another mechanism by which intestinal lipids suppress food intake is by promoting synthesis of fatty acid ethanol amides (FAEs). In particular, intestinal OEA synthesized from dietary lipid after a meal is known to induce satiety [14; 15; 30; 52]. Recent studies indicate that BAs stabilize and activate the FAE-producing enzyme NAPE-PLD [12; 53]. To investigate the involvement of pythocholic acid in NAPE-PLD and FAE production, we measured 3 FAE species–OEA, AEA and DHEA–in jejunal epithelia. Mice were given the pythocholic acid or DCA once a day for two days. We collected jejunal epithelia after food intake and gastric emptying experiment (Figure 8A). Jejunal OEA, AEA, and DHEA levels after refeeding were robustly increased in pythocholic acid-treated mice relative to vehicle or DCA (Figure 8B-D).

**Figure 8.**
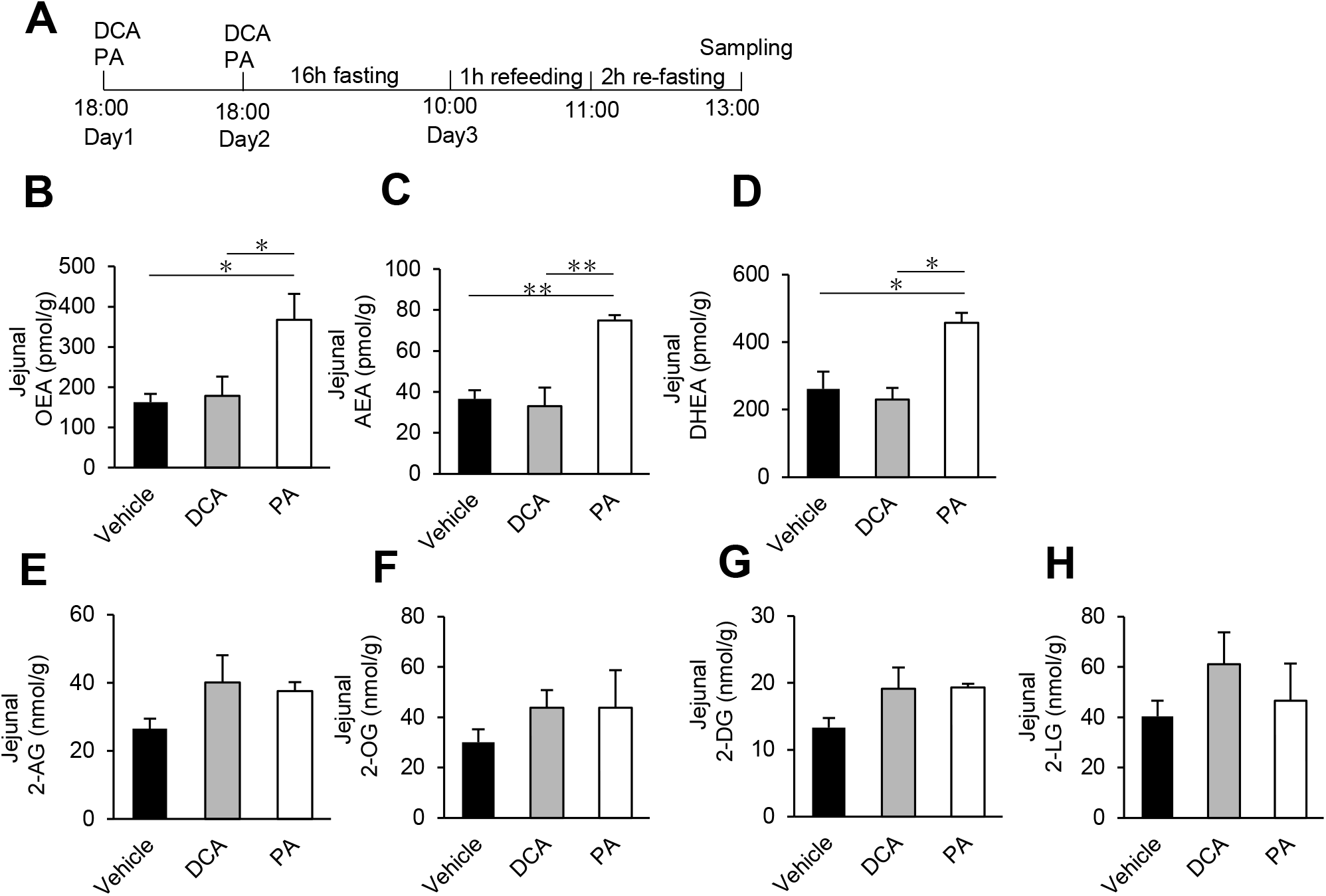
Effects of DCA and PA on fatty acid ethanolamides and cannabinoids in jejunal intestine after refeeding. (A) experimental schedule, Effect of DCA and PA after refeeding on the levels of (B) OEA, (C) AEA, (D) DHEA, (E) 2-AG, (F) 2-OG, (G) 2-DG and (H) 2-LG. (Vehicle; n=5, DCA; n=5, PA; n=3). \**p*<0.05, \*\**p*<0.01 (one-way repeated ANOVA).

We measured several other bioactive lipids, 2-AG, 2-OG, 2-DG and 2-LG, which are synthesized from phospholipids by diacylglycerol lipase (DGL). We found no differences in jejunal 2-AG, 2-OG, 2-DG or 2-LG between treatment groups (Figure 8E-H). These data indicate that pythocholic acid promotes NAPE-PLD-dependent, but not DGL-dependent, bioactive lipid synthesis in the jejunum.

## 4. DISCUSSION

In this study, we successfully isolated pythocholic acid from pythons for experimental use and tested the effect of pythocholic acid on food intake in mice. Our data showed that pythocholic acid treatment decreased ad libitum food intake and increased FAEs, especially OEA in the jejunum. In addition, the anorexigenic effect of pythocholic acid was independent of lipid absorption and gastric emptying. Our findings indicate that pythocholic acid is a novel tool to promote intestinal OEA and suppress food intake.

Feeding promotes jejunal OEA formation in chow diet fed-lean rodents [11; 15; 30; 31]. There are three essential steps to produce and maintain OEA: (i) production of oleic acid-containing NAPEs, (ii) NAPE-PLD activation to produce OEA from oleic acid-containing NAPEs and (iii) inhibition of fatty acid amide hydrolase (FAAH) that hydrolyzes OEA [16; 30]. Among these biological events that maintain OEA, BAs are known to play a role in NAPE-PLD activation. Hydroxyl groups at carbon 3α and 12α– in BAs bind the substrate-binding pocket of the NAPE-PLD dimer, which reinforces the enzyme’s stability and activity [12]. Kinetic experiments demonstrated that DCA (3α, 12α) increases NAPE-PLD activity better than chenodeoxycholic acid, another dihydroxy BA (3α, 7α) [37]. Our data showed that the tri-hydroxylated BA (3α, 12α, 16α), pythocholic acid increased jejunal OEA levels much higher than DCA in mice. Also, pythocholic acid increased two other FAEs–AEA and DHEA–which are also formed by NAPE-PLD activation. But we found no effect of pythocholic acid on other bioactive lipids levels such as 2-AG, 2-OG, 2-DG and 2-LG, which are synthesized by monoacylglycerol lipase. These findings suggest that the specific structure of pythocholic acid activates NAPE-PLD in the intestine. However, further studies would be needed to show the direct binding between NAPE-PLD and pythocholic acid.

Though the mechanism is not known, OEA mobilization is disrupted in diet-induced obese rodents [54]. On the other hand, OEA treatment decreases food intake and body weight in diet-induced obese rodents [21]. This evidence suggests that impaired OEA synthesis contributes to obesity. Examining the effect of pythocholic acid on food intake, body weight gain and glucose homeostasis in diet-induced obese rodents is a future direction of this work.

Another possible pathway to regulate food intake by BAs is TGR5-mediated signaling in the brain. Feeding or oral gavage of BAs allows access of BAs into the hypothalamus, which decreases the food intake [10]. Intracerebroventricular injection of TGR5 specific agonist, -(2-chlorophenyl)-N-(4-chlorophenyl)-N,5-dimethyl-4-isoxazolecarboxamid (CCDC) decreases food intake [4]. Deletion of TGR5 in AgRP neuron cancels the anorexigenic effect of TGR5 agonist INT-777 [10]. Thus, TGR5 signaling in the central nervous system is an alternative pathway for food intake regulation by BAs. Further studies to evaluate the transport of pythocholic acid into the brain after oral gavage and its effects on TGR5 will be of interest to fully understand the anorexigenic mechanism(s) of pythocholic acid.

Pythocholic acid increased not only OEA but also AEA in the jejunum. AEA is a natural endogenous ligand for cannabinoid (CB) 1 receptor which induces feeding behavior via hypothalamic signaling [52; 55; 56]. The role of intestinal cannabinoid signaling on food intake regulation has also been established. Starvation promotes intestinal AEA levels, and feeding decreases AEA levels [57]. Peripheral AEA treatment increases food intake, and the CB1 receptor antagonist SR151716A decreases food intake in rodents [57]. Another study shows that mice consuming a western style diet display hyperphagia with increased levels of 2-AG and AEA in the jejunum, and this can be blocked by CB1 receptor antagonism [40]. Thus the anorexigenic effect of intestinal OEA and orexigenic effect of AEA are inversely correlated. Our data showed both OEA and AEA were increased in the intestine by pythocholic acid treatment. Gomez and colleagues reported that OEA treatment blocks the AEA-induced hyperphagia [57]. In agreement with this report, pythocholic acid decreased food intake, suggesting that the anorexigenic effect of elevated OEA outweighed the orexigenic effect of AEA.

In conclusion, our findings reveal an anorexigenic effect of the python-derived 16α-hydroxylated bile acid, pythocholic acid. Results in this study indicate that modification of bile acid, particularly 16α-hydroxylation or pythocholic acid treatment may be a novel strategy for obesity and metabolic disease treatment.

## Abbreviations

BA: bile acid
CA: cholic acid
DCA: deoxycholic acid
CDCA: chenodeoxycholic acid
ALT: alanine transaminase
AST: aspartate transaminase
FFA: free fatty acids
TG: Triglyceride
NAPE-PLD: N-acyl phosphatidylethanolamine phospholipase D
FAEs: fatty acid ethanolamides
OEA: oleoylethanolamide
AEA: arachidonoylethanolamide (also known as anandamide)
DHEA: docosahexaenoylethanolamide
2-AG: 2-arachidonylglycerol
2-OG: 2-oleoylglycerol
2-DG: 2-decanoyl-rac-glycerol
2-LG: 2-linoleoylglycerol

## ACKNOWLEDGMENTS

We thank members of the Haeusler lab, Dr. Domenico Accili (Columbia University), Dr. Kimberly Lackey, Dr. Stephen Secor (University of Alabama), Mr. Dustin Leahy (Piedthonidae Exotics), Dr. Reiko Toyama (NICHD) and Dr. Jergen Wess (NIDDK), and Arclev Academia Strategists Network (AASN) members for helpful discussions, suggestions, and/or providing materials. I got this research idea from my kid’s pet snake (Albino ball python), so I thank Reiri and Akira Higuchi for giving research ideas and caring for their pet snake.

## Contributors

S.H., and R.A.H conceived and designed research; S.H., C.W. A.K., performed experiments; S.H., N.V.D. and A.K. analyzed data; S.H. and R.A.H drafted the manuscript; all authors edited the manuscript.

## Funding

Funding was from the National Institutes of Health R01DK115825 to RAH, the American Diabetes Association 7-20-IBS-130 to RAH, and from NIH-funded center grants P30DK026687 and P30DK132710.

## Competing interest

The authors disclose no conflicts.

## Notes

### Competing Interest Statement

The authors have declared no competing interest.

